# Assessing the impact of free-roaming dog population management through systems modelling

**DOI:** 10.1101/2021.12.02.470957

**Authors:** Lauren M. Smith, Rupert J. Quinnell, Conor Goold, Alex M. Munteanu, Sabine Hartmann, Lisa M. Collins

**Affiliations:** School of Biology, Faculty of Biological Sciences, University of Leeds, Leeds, UK, LS2 9JT; VIER PFOTEN International, Vienna, Austria

## Abstract

Free-roaming dogs can present significant challenges to public health, wildlife conservation, and livestock production. Their own welfare may also be a concern, as free-roaming dogs can experience poor health and welfare. Dog population management is widely conducted to mitigate these issues. To ensure efficient use of resources, it is critical that effective, cost-efficient, and high-welfare strategies are identified. The dog population comprises distinct subpopulations characterised by their restriction status and level of ownership, but the assessment of dog population management often fails to consider the impact of the interaction between subpopulations on management success. We present a system dynamics model that incorporates an interactive and dynamic system of dog subpopulations. We identify that methods incorporating both fertility control and responsible ownership interventions (a reduction in abandonment and an increase in shelter adoptions) have the greatest potential to reduce free-roaming dog population sizes over longer periods of time, whilst being cost-effective and improving overall welfare. We suggest that future management should be applied at high levels of coverage and should target all sources of population increase, such as abandonment, births, and free-roaming owned dogs, to ensure effective and cost-efficient reduction in free-roaming dog numbers.

## Introduction

There are between 700 million and one billion dogs globally, comprising owned and unowned dogs that live in shelters, homes, or freely in urban and rural environments ^[1,2]^. Where dogs freely roam and exist in high densities, they can present significant challenges to public health ^[3,4]^, wildlife conservation ^[5]^, and livestock production ^[6,7]^. Free-roaming dogs can also experience conditions leading to poor health and welfare states ^[8,9]^. Population management is conducted to control free-roaming dog population size and, depending on the approach taken, to improve dog health and welfare and mitigate public health and conservation problems ^[10,11]^.

Dog populations can be divided into four subpopulations depending on their relationship with humans (owned or unowned) and their restriction status (restricted or free-roaming). The free-roaming dog population (i.e. street dogs, including owned and unowned dogs) is of most interest for dog population management ^[12]^. Dog subpopulations are interactive and dynamic – individuals can move between different subpopulations through time (Figure 1). Throughout its life, a dog could spend time in multiple states – for example, living as an unowned stray dog, being caught and moved into shelter accommodation, before being adopted into a home.

**Figure 1.**
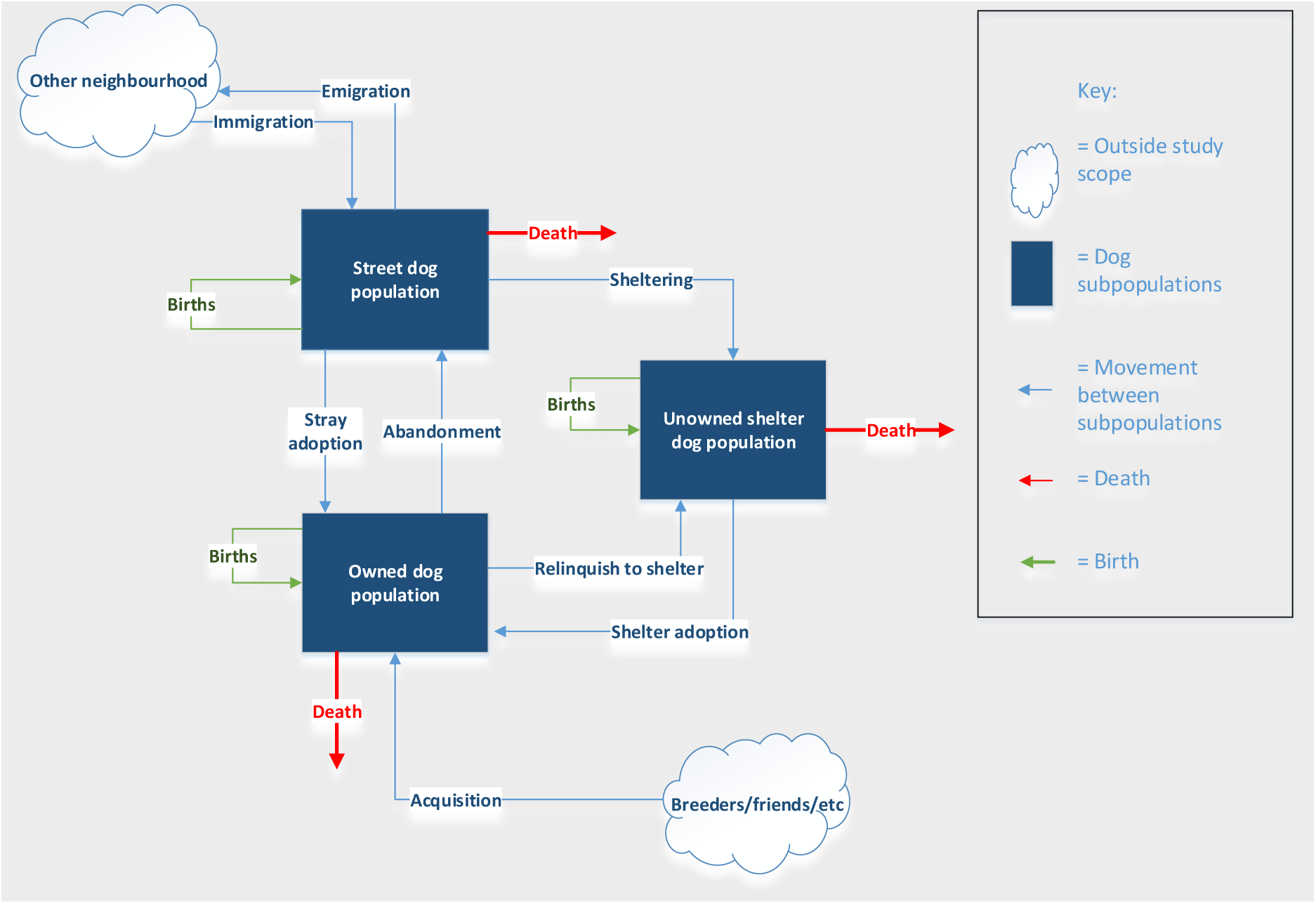
Process of dog subpopulation increase/decrease, including movement of dogs between different dog subpopulations.

Dog population management can involve reproductive control (e.g. catch-neuter-release; CNR), sheltering, culling, and responsible ownership campaigns ^[10,12]^. The free-roaming dog population size can be reduced by decreasing recruitment into (e.g. births and immigration) and increasing removal from (e.g. mortality and emigration) the population. As the dog population consists of interacting subpopulations, migration not only occurs between free-roaming dog populations in different geographic locations, but also between the subpopulations within a location, providing additional sources of recruitment. In terms of their effect on recruitment and removal, both culling and sheltering aim to increase rates of removal from the free-roaming dog population. Both CNR and responsible ownership campaigns aim to decrease rates of recruitment to the free-roaming dog population: CNR through decreased birth rates; and responsible ownership campaigns, depending on the approach, through the reduced abandonment of owned dogs.

Modelling methods can aid in planning, implementing, and evaluating dog population management strategies. System dynamics modelling allows complex and interactive systems to be explained and the impact of interventions on model behaviour to be evaluated ^[13,14]^. Dog population dynamics have been investigated with both system dynamics approaches ^[15–23]^ and agent-based models ^[24,25]^, assessing the effects of management strategies on population size ^[15,21,22,25,26]^, euthanasia rates ^[23]^, and disease dynamics ^[19,20]^. Most previous studies have modelled dynamics within a single subset of the population (e.g. owned or free-roaming). Few have modelled the interactions between these subpopulations. Modelling these interactions could provide greater insight into the effectiveness of dog population management interventions in reducing dog population size. All population management methods require the investment of resources (e.g. staff, facilities, and equipment), but few previous models have considered cost-effectiveness ^[16,22,23]^.

The aim of this study was to compare the impact of different dog population management methods on the sizes of the different dog subpopulations, the staff-resources required, and the welfare costs of each intervention using a system dynamics modelling approach. The management methods under investigation included sheltering, culling, CNR, and responsible ownership. To reflect that management practices often involve combinations of management methods ^[27]^, we also investigate combined CNR and sheltering, and combined CNR and responsible ownership.

## Methods

### Model description

The system dynamics model divided an urban dog population into the following subpopulations: (i) street dogs (owned and unowned free-roaming), (ii) shelter dogs (unowned restricted), and (iii) owned dogs (owned restricted) (Figure 1). The subpopulations change in size by individuals flowing between the different subpopulations or from flows extrinsically modelled (e.g. acquisition of dogs from breeders and friends).

Ordinary differential equations were used to describe the dog population dynamics. The models were written in R version 3.6.1 ^[28]^, and numerically solved using the Runge-Kutta fourth order integration scheme with a 0.01 step sizes using the package “deSolve” ^[29,30]^. For the baseline model, Equations 1, 2, and 3 were used to describe the rates of change of dog subpopulations in the absence of management.

*Equation 1. Baseline street dog population (S).*

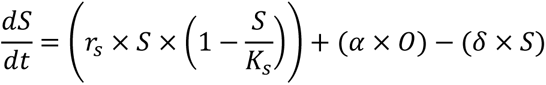

*Equation 2. Baseline shelter dog population (H).*

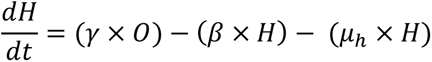

*Equation 3. Baseline owned dog population (O).*

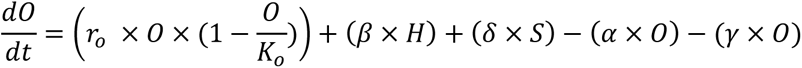

In the baseline model, the street dog population (Equation 1) increases through the street dog intrinsic growth rate (*r_s_*), and abandonment of dogs from the owned dog population (*α*) and decreases through adoption to the owned dog population (*δ*). The intrinsic growth rate is the sum of the effects of births, deaths, immigration, and emigration, which are not modelled separately. In this model, the growth rate of the street dog population is reduced depending on the population size in relation to the carrying capacity, through the logistic equation (*r_real_* = *r_max_*(1-*S*/*K_s_*)) ^[31]^. In the baseline simulation, the street dog population rises over time, until it stabilises at an equilibrium size.

The shelter dog population (Equation 2) increases through relinquishment of owned dogs (*γ*) and decreases through the adoption of shelter dogs to the owned dog population (*β*). There is no carrying capacity for the shelter dog population (i.e. it is assumed that more housing would be created as the population increases), allowing calculation of resources required to house shelter dogs.

The owned dog population (Equation 3) increases through the owned dog growth rate (*r_o_*), adoption of shelter dogs (*β*), and adoption of street dogs (*δ*); and decreases through abandonment (*α*) and relinquishment (*γ*) of owned dogs to the shelter dog population. The growth rate of the owned dog population (*r_o_*) combines the birth, death, and acquisition rates from sources other than the street or shelters (e.g. breeders, friends) and was modelled as density dependent by the limit to growth logistic formula (1-O/K_o_).

### Parameter estimates

Detailed descriptions of parameter estimates are provided in the supplementary information. The simulated environment was based on the city of Lviv, Ukraine. This city has an area of 182km^2^ and a human population size of 717,803. Parameters were estimated from literature, where possible, and converted to monthly rates (Table 1). Initial sizes of the dog populations were estimated for the baseline simulation, based on our previous research in Lviv ^[32]^. The carrying capacity depends on the availability of resources (i.e. food, shelter, water, and human attitudes and behaviour ^[33]^) and is challenging to estimate. We assumed the initial street and the owned dog populations were at carrying capacity. Initial population sizes for simulations including interventions were determined by the equilibrium population sizes from the baseline simulation (i.e. the stable population size, the points at which the populations were no longer increasing/decreasing).

**Table 1.** Parameter description, parameter value, and minimum and maximum values used in the sensitivity analysis for the systems model. Rates are per month.

| Parameter | Description | Value | Min | Max | Unit | Reference |
| --- | --- | --- | --- | --- | --- | --- |
| $S$ | Initial street dog population for baseline simulation | 20,000 | NA | NA | Dogs | Extrapolated from <sup>[32]</sup> |
| $H$ | Initial shelter dog population for baseline simulation | 3750 | NA | NA | Dogs | Estimated from Lviv local shelter data |
| $O$ | Initial owned dog population for baseline simulation | 100,492 | NA | NA | Dogs | Extrapolated from <sup>[32]</sup> |
| $K_s$ | Carrying capacity of street dog population | 20,000 | 15,000 | 25,000 | Dogs | Extrapolated from <sup>[32]</sup> |
| $K_o$ | Carrying capacity of owned dog population | 100,492 | 90,000 | 110,000 | Dogs | Extrapolated from <sup>[32]</sup> |
| $\alpha$ | Abandonment rate of owned dogs to street population | 0.003 | 0.001 | 0.006 | 1/month | <sup>[35]</sup> |
| $\beta$ | Adoption rate of shelter dogs to owned population | 0.025 | 0.015 | 0.035 | 1/month | Estimated from Lviv local shelter data |
| $r_s$ | Maximum growth rate of street dog population | 0.03 | 0.01 | 0.1 | 1/month | <sup>[17,19,37]</sup> |
| $r_o$ | Maximum growth rate of owned dog population | 0.07 | 0.0357 | 0.1125 | 1/month | Assumption |
| $\gamma$ | Relinquishment rate of owned dogs to shelter population | 0.0007 | 0.0003 | 0.001 | 1/month | <sup>[36]</sup> |
| $\delta$ | Adoption rate of street dogs to owned population | 0.007 | 0.0035 | 0.0105 | 1/month | Estimated from Lviv local shelter data |
| $\mu_h$ | Death rate of shelter dogs | 0.008 | 0.005 | 0.02 | 1/month | Assumption |
| $\mu_s$ | Minimum death rate of neutered street dogs | 0.02 | NA | NA | 1/month | [37–40] |

Estimating the rate at which owned dogs are abandoned (*α)* is difficult, as abandonment rates are often reported per dog-owning lifetime ^[32,34]^ and owners are likely to under-report abandonment of dogs. We derived the abandonment rate from Fielding & Plumridge (2005) ^[35]^ and the owned dog relinquishment rate (*γ*) from New *et al.,* (2004) ^[36]^. We estimated shelter (*β*) and street adoption rates (*δ*) from shelter data in Lviv. We set the maximum intrinsic growth rate for the street dogs (*r_s_*) at 0.03 per month, similar to that reported in literature ^[17,19,37]^. We assumed that demand for dogs was met quickly through a supply of dogs from births, breeders and friends and set a higher growth rate for the owned dog population (*r_o_*) at 0.07 per month.

We assumed shelters operated with a “no-kill” policy (i.e. dogs were not killed in shelters as part of population management) and included a shelter dog death rate (*μ_h_*) of 0.008 per month to incorporate deaths due to behavioural problems, health problems and natural mortality. We modelled neutered street dog death rate (*μ_s_*) explicitly for the CNR intervention at a minimum death rate of 0.02 per month ^[37–40]^.

### Interventions

Six intervention scenarios were modelled (Table 2): sheltering; culling; CNR; responsible ownership; combined CNR and responsible ownership; and combined CNR and sheltering, representing interventions feasibly applied and often reported ^[27]^. Table 2 outlines the equations describing each intervention. To simulate a sheltering intervention, a proportion of the street dog population was removed and added to the shelter dog population at sheltering rate (σ). In culling interventions, a proportion of the street dog population was removed through culling (χ).

**Table 2.**
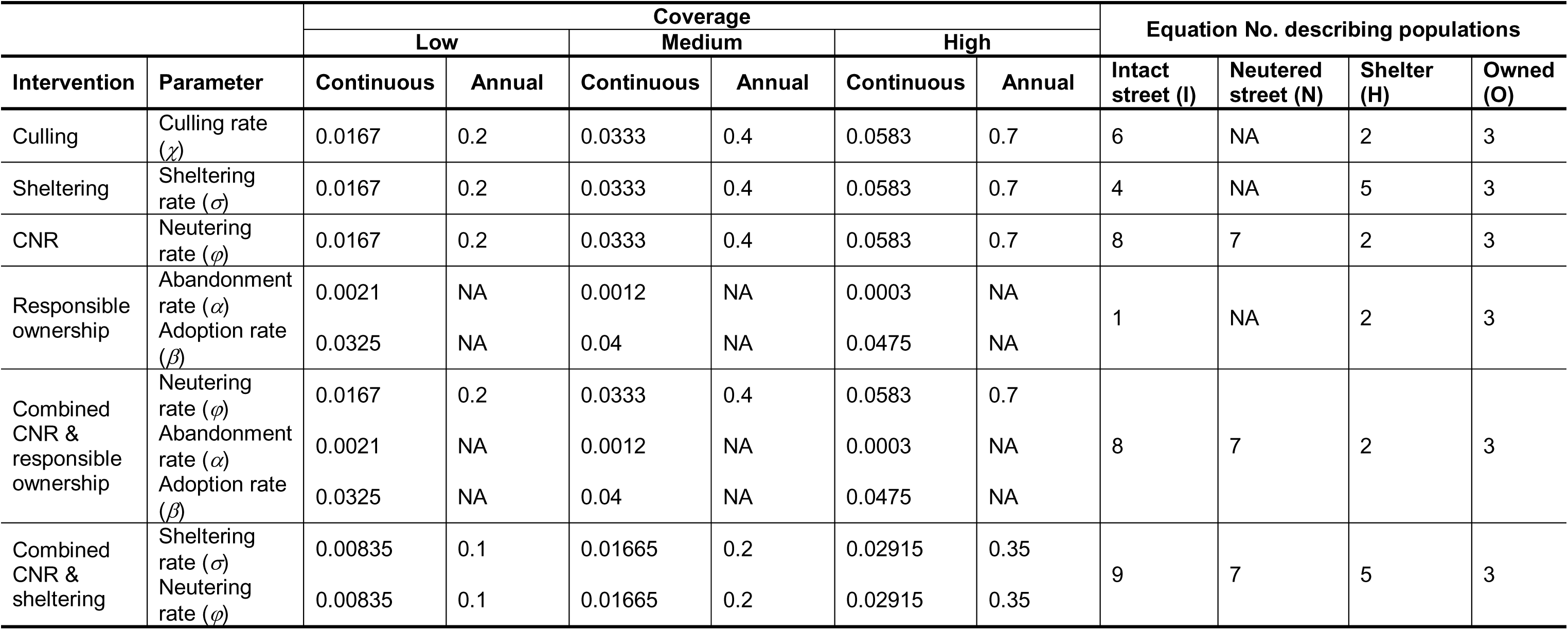
Description of intervention parameters and coverages for simulations applied at continuous and annual periodicities.

|  |  | Coverage |  |  |  |  |  | Equation No. describing populations |  |  |  |
| --- | --- | --- | --- | --- | --- | --- | --- | --- | --- | --- | --- |
|  |  | Low |  | Medium |  | High |  |  |  |  |  |
| Intervention | Parameter | Continuous | Annual | Continuous | Annual | Continuous | Annual | Intact street (I) | Neutered street (N) | Shelter (H) | Owned (O) |
| Culling | Culling rate ($\chi$) | 0.0167 | 0.2 | 0.0333 | 0.4 | 0.0583 | 0.7 | 6 | NA | 2 | 3 |
| Sheltering | Sheltering rate ($\sigma$) | 0.0167 | 0.2 | 0.0333 | 0.4 | 0.0583 | 0.7 | 4 | NA | 5 | 3 |
| CNR | Neutering rate ($\varphi$) | 0.0167 | 0.2 | 0.0333 | 0.4 | 0.0583 | 0.7 | 8 | 7 | 2 | 3 |
| Responsible ownership | Abandonment rate ($\alpha$) | 0.0021 | NA | 0.0012 | NA | 0.0003 | NA | 1 | NA | 2 | 3 |
| | Adoption rate ($\beta$) | 0.0325 | NA | 0.04 | NA | 0.0475 | NA | | | | |
| Combined CNR & responsible ownership | Neutering rate ($\varphi$) | 0.0167 | 0.2 | 0.0333 | 0.4 | 0.0583 | 0.7 | 8 | 7 | 2 | 3 |
| | Abandonment rate ($\alpha$) | 0.0021 | NA | 0.0012 | NA | 0.0003 | NA | | | | |
| | Adoption rate ($\beta$) | 0.0325 | NA | 0.04 | NA | 0.0475 | NA | | | | |
| Combined CNR & sheltering | Sheltering rate ($\sigma$) | 0.00835 | 0.1 | 0.01665 | 0.2 | 0.02915 | 0.35 | 9 | 7 | 5 | 3 |
| | Neutering rate ($\varphi$) | 0.00835 | 0.1 | 0.01665 | 0.2 | 0.02915 | 0.35 | | | | |

*Equation 4. Street dog population with sheltering intervention*

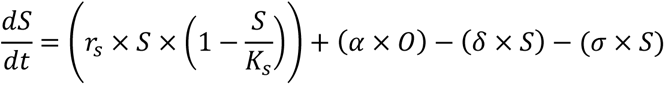

*Equation 5. Shelter dog population with sheltering intervention.*

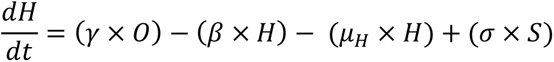

*Equation 6. Street dog population with a culling intervention.*

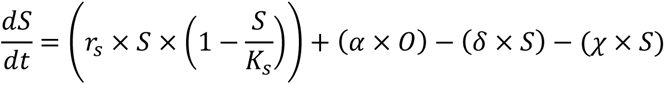

To simulate a CNR intervention, an additional subpopulation was added to the system (Equation 7): (iv) the neutered street dog population (*N*; neutered, free-roaming). In this simulation, a proportion of the intact (*I*) street dog population was removed and added to the neutered street dog population. A neutering rate (*φ*) was added to the differential equations describing the intact street and the neutered street dog populations. Neutering was assumed to be lifelong (e.g. gonadectomy); a neutered street dog could not re-enter the intact street dog subpopulation. Neutered street dogs were removed from the population through the density dependent neutered dog death rate (*μ_n_*); death rate increased when the population was closer to the carrying capacity. The death rate was a non-linear function of population size and carrying capacity modelled using a table lookup function (Figure S1).

*Equation 7. Neutered street dog population stock*

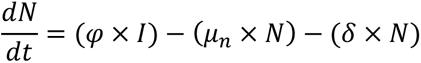

*Equation 8. Intact street dog population with neutering intervention.*

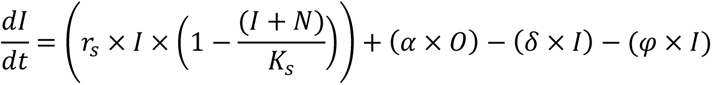

To simulate a responsible ownership intervention, the baseline model was applied with decreased rate of abandonment (*α*) and increased rate of shelter adoption (*β*). To simulate combined CNR and responsible ownership, a proportion of the intact street dog population was removed through the neutering rate (*φ*), abandonments decreased (*α*) and shelter adoptions increased (*β*). In combined CNR and sheltering interventions, a proportion of the intact street dog population (*I*) was removed through neutering (*φ*) and added to the neutered street dog population (*N*), and a proportion was removed through sheltering (σ) and added to the shelter dog population (*H*).

*Equation 9. Intact street dog population with combined CNR and sheltering interventions.*

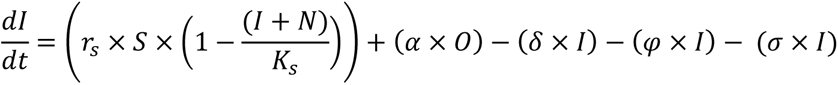

### Intervention length, periodicity, and coverage

All simulations were run for 70 years. Interventions were applied for two lengths of time: (i) the full 70-year duration of the simulation; and (ii) a five-year period followed by no further intervention, to simulate a single period of investment in population management. In each of these simulations, we modelled the interventions as (i) continuous (i.e. a constant rate of e.g. neutering) and (ii) annual (i.e. intervention applied once per year). Interventions were run at low, medium, and high coverages (Table 2). Intervention coverage refers to the proportion of dogs that are culled/neutered/sheltered per year (i.e. 20%, 40% and 70% annually) and, for responsible ownership interventions, the decrease in abandonment rate and increase in the adoption rate of shelter dogs (30%, 60% and 90% increase/decrease from baseline values). To model a low (20%), medium (40%) and high (70%) proportion of street dogs caught, but where half of the dogs were sheltered and half were neutered-and-returned, combined CNR and sheltering interventions were simulated at half-coverage (e.g. intervention rate of 0.7 was simulated by 0.35 neutered and 0.35 sheltered). For continuous interventions, sheltering (*σ*), culling (*χ*), and CNR (*φ*) were applied continuously during the length of the intervention. For annual interventions, *σ, χ*, and *φ* were applied to the ordinary differential equations using a forcing function applied at 12-month intervals. In simulations that included responsible ownership interventions, the decrease in owned dog abandonment (*α*) and the increase in shelter adoption (*β*) was assumed instantaneous and continuous (i.e. rates did not change throughout the intervention).

### Model outputs

The primary outcome of interest was the impact of interventions on street dog population size. For interventions applied for the duration of the simulation, we calculated: (i) equilibrium population size for each population; (ii) percent decrease in street dog population; (iii) costs of intervention in terms of staff-time; and (iv) an overall welfare score. For interventions applied for a five-year period, we also calculated: (v) minimum street dog population size and percent reduction from initial population size; and (vi) the length of time between the end of the intervention and time-point at which the street dog population reached above 20,000 dogs.

The costs of population management interventions vary by country (e.g. staff salaries vary between countries) and by the method of application (e.g. method of culling, or resources provided in a shelter). To enable a comparison of the resources required for each intervention, the staff time (staff working-months) required to achieve the intervention coverage was calculated. While this does not incorporate the full costs of an intervention, as equipment (e.g. surgical equipment), advertising campaigns, travel costs for the animal care team, and facilities (e.g. clinic or shelter costs) are not included, it can be used as a proxy for intervention cost. Using data provided from VIER PFOTEN International, we estimated the average number of staff required to catch and neuter the street dog population and to house the shelter dog population in each intervention, using this data as a proxy for catching and sheltering/culling. The number of dogs that can be cared for per shelter staff varies by shelter. To account for this, we estimated two staff-to-dog ratios (low and high). Table 3 describes the staff requirements for the different interventions.

**Table 3.** Staff required for interventions and the number of dogs processed per staff per day.

| Staff type | Baseline | CNR | Sheltering | Culling | Responsible ownership | Dogs/Staff/Day | Relative salary |
| --- | --- | --- | --- | --- | --- | --- | --- |
| Veterinarian | No | Yes | No | No | No | 6 | 1.00 |
| Veterinary nurse | No | Yes | No | No | No | 8 | 0.65 |
| Dog catchers | No | Yes | Yes | Yes | No | 13 | 0.56 |
| Kennel staff | Yes | Yes | Yes | Yes | Yes | 13 (low)<br>120 (high) | 0.50 |
| Campaigner | No | No | No | No | Yes | NA | 0.75 |

Using the projected population sizes, the staff time required for each staff type (e.g. number of veterinarian-months of work required) was calculated for each intervention. Relative salaries for the different staff types were estimated (Table 3). The relative salaries were used to calculate the cost of the interventions by:

[*staff time required* x *relative salary* ] x €20,000.

Where €20,000 was the estimated annual salary of a European veterinarian, allowing relative staff-time costs to be compared between the different interventions. Average annual costs were reported.

To provide overall welfare scores for each of the interventions, we apply the estimated welfare scores on a one to five scale, for each of the dog subpopulations, as described by Hogasen *et al.* (2013) ^[22]^. This scale is based on the Five Freedoms (freedom from hunger and thirst; freedom from discomfort; freedom from pain, injury, or disease; freedom to express normal behaviour; freedom from fear and distress ^[41,42]^) and was calculated using expert opinions from 60 veterinarians in Italy. The scores were weighted by the participants’ self-reported knowledge of different dog subpopulations, which resulted in the following scores: 2.8 for shelter dogs (W_H_); 3.5 for owned dogs (W_O_); 3.1 for neutered street dogs (W_N_); and 2.3 for intact street dogs (W_I_).

Using these estimated welfare scores, we calculated an average welfare score for the total dog population based on the model’s projected population sizes for each subpopulation (Equation 10). For interventions running for the duration of the simulation, the welfare score was calculated at the time point (*t*) when the population reached an equilibrium size. For interventions running for five years, the welfare score was calculated at the end of the five - year intervention. The percentage change in welfare scores from the baseline simulation were reported.

Equation 10

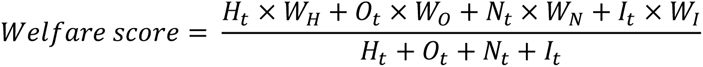

### Model validation and sensitivity analysis

A global sensitivity analysis was conducted on all parameters described in the baseline simulation and all interventions applied continuously, at high coverage, for the full duration of the simulation. A Latin square design algorithm was used in package “FME” ^[43]^ to sample the parameters within their range of values (Table 1). For the global sensitivity analysis on interventions, all parameter values were varied, apart from the parameters involved in the intervention (e.g. culling, neutering, abandonment rates). The effects of altering individual parameters (local sensitivity analysis) on the population equilibrium was also examined for the baseline simulation using the Latin square design algorithm to sample each parameter, individually, within their range of values. Sensitivity analyses were run for 100 simulations over 50 years solved with 0.01 step sizes.

## Results

### Model validation and sensitivity analysis in baseline simulation

The baseline simulation resulted in equilibrium sizes of 23,651, 2086, and 98,358 for the street, shelter, and owned populations respectively. The population sizes settle at an equilibrium close, but not equal to the carrying capacities for the street or owned populations due to the flows of abandonment (*α*), relinquishment (*γ*), and adoption from streets (*δ*) and shelters (*β*). In the global sensitivity analysis, for the baseline simulation, the simulated populations stabilised at an equilibrium between (A) 14,267 and 35,011 street dogs; (B) 596 and 4140 shelter dogs; (C) 84,737 and 109,236 owned dogs (Figure S2). Results of the local sensitivity analysis are presented in Figures S3 and S4.

### Impacts of interventions applied for full duration of simulation (70 years)

Table 4 outlines the impact of interventions applied for the full duration of the simulation. All interventions reduced the street dog population size (Figure 2). Combined CNR and responsible ownership had the greatest reduction in street dog equilibrium population size at high (annual = 89%, continuous = 90% reduction) and medium (annual = 55%, continuous = 57% reduction) coverages. Culling most greatly reduced the population at low coverages (annual = 29%, continuous = 31%), although both sheltering and combined CNR and responsible ownership had similar effects. At all coverages, responsible ownership and CNR applied alone had only small effects on the street dog population, whereas culling and sheltering both had large effects. Combined CNR and sheltering was more effective than CNR, but less effective than sheltering applied alone.

**Figure 2.**
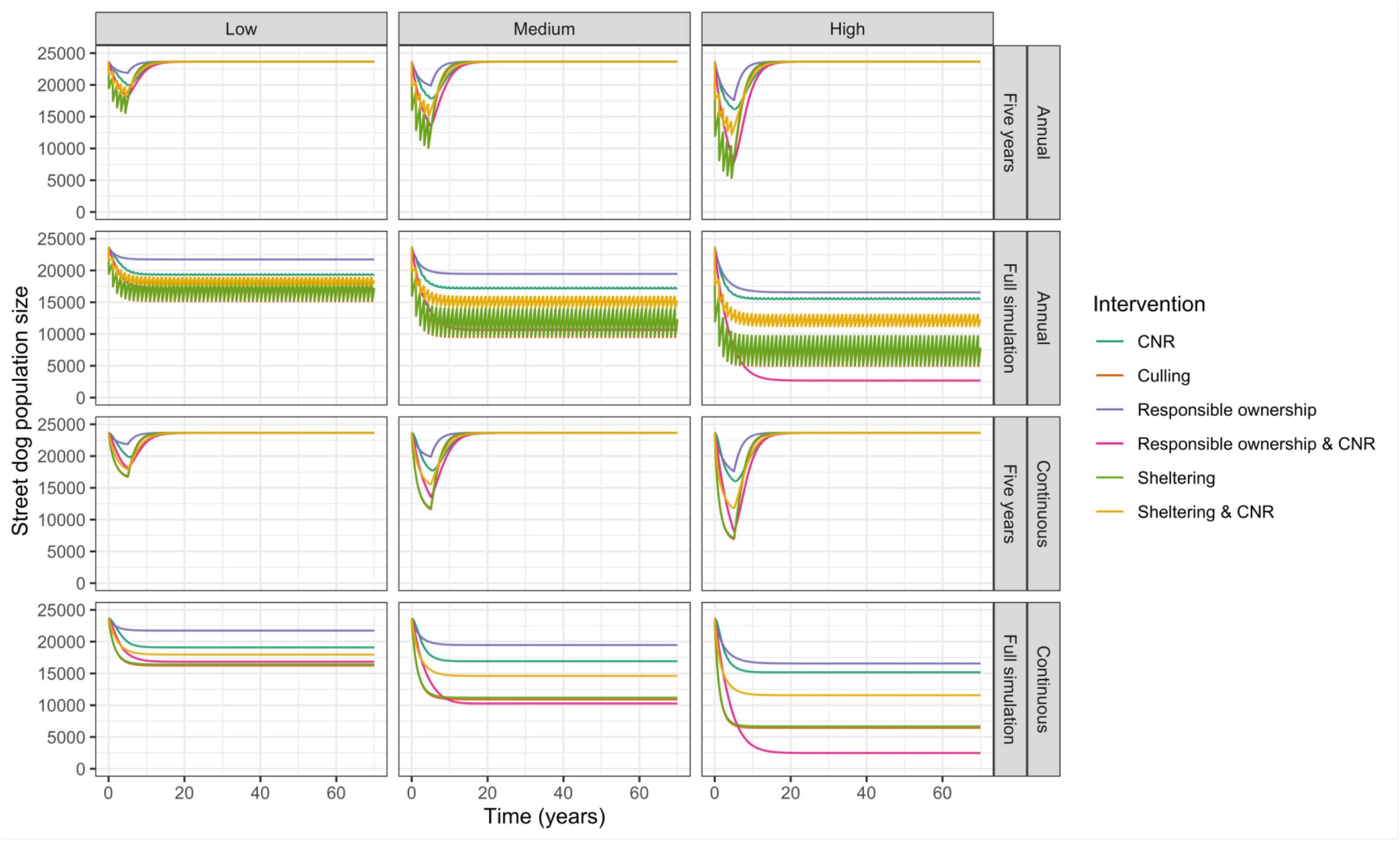
Impact of interventions on street dog population for simulations run with continuously applied control and five-year period of intervention, when interventions are applied annually and continuously at low, medium, and high coverages.

**Table 4.** Impact of interventions on street (S), shelter (H) and owned (O) dog population equilibrium levels and percent change from baseline population equilibrium levels for interventions applied for the duration of the simulation at low, medium, and high coverages. The intervention with the greatest reduction in street dog population size for each periodicity-coverage combination is highlighted in grey. The greatest increase in welfare scores and the lowest average annual costs for each periodicity-coverage combination are highlighted in bold.

| Periodicity | Coverage | Intervention | Equilibrium population size |  |  | Percentage population change (%) |  |  | Welfare score and % change | Average annual cost (€) |  |
| --- | --- | --- | --- | --- | --- | --- | --- | --- | --- | --- | --- |
|  |  |  | S | H | O | S | H | O |  | Shelter staff-to-dog ratio |  |
|  |  |  |  |  |  |  |  |  |  | High | Low |
| NA | NA | Baseline | 23651 | 2086 | 98358 | 0 | 0 | 0 | 3.26 (NA) | 119,559 | 1,103,620 |
| Annual* | Low | CNR | 19364 | 2065 | 97358 | -18 | -1 | -1 | 3.33 (2.13%) | 164,965 | 1,179,519 |
|  |  | CNR & responsible ownership | 17130 | 1702 | 98454 | -28 | -18 | 0 | **3.35 (2.84%)** | 159,548 | **1,002,944** |
|  |  | 0.5 CNR & 0.5 sheltering | 18316 | 5598 | 98720 | -23 | 168 | 0 | 3.31 (1.54%) | 351,178 | 3,048,653 |
|  |  | **Culling** | **16690** | 2072 | 97671 | **-29** | -1 | -1 | 3.31 (1.62%) | **131,028** | 1,142,971 |
|  |  | Sheltering | 16882 | 10063 | 100428 | -29 | 382 | 2 | 3.28 (0.76%) | 588,204 | 5,367,903 |
|  | Medium | CNR | 17232 | 2054 | 96854 | -27 | -2 | -2 | 3.37 (3.27%) | 181,908 | 1,191,651 |
|  |  | **CNR & responsible ownership** | **10714** | 1445 | 99117 | **-55** | -31 | 1 | **3.42 (4.77%)** | 156,501 | **877,309** |
|  |  | 0.5 CNR & 0.5 sheltering | 15113 | 6664 | 98587 | -36 | 219 | 0 | 3.34 (2.60%) | 429,451 | 3,687,716 |
|  |  |  | S | H | O | S | H | O |  | Shelter staff-to-dog ratio |  |
|  |  |  |  |  |  |  |  |  |  | High | Low |
|  |  | Culling | 11630 | 2061 | 97155 | -51 | -1 | -1 | 3.36 (2.99%) | **134,410** | 1,146,945 |
|  |  | Sheltering | 11898 | 13217 | 101004 | -50 | 533 | 3 | 3.31 (1.62%) | 784,217 | 7,143,262 |
|  | High | CNR | 15570 | 2047 | 96523 | -34 | -2 | -2 | 3.39 (4.07%) | 196,557 | 1,203,011 |
|  |  | **CNR & responsible ownership** | **2682** | 1261 | 100007 | **-89** | -40 | 2 | **3.47 (6.58%)** | 139,026 | **770,623** |
|  |  | 0.5 CNR & 0.5 sheltering | 12187 | 7046 | 98364 | -48 | 238 | 0 | 3.37 (3.48%) | 466,462 | 3,961,066 |
|  |  | Culling | 7231 | 2051 | 96698 | -69 | -2 | -2 | 3.40 (4.33%) | **136,569** | 1,144,708 |
|  |  | Sheltering | 7474 | 14015 | 100849 | -68 | 572 | 3 | 3.35 (2.65%) | 851,283 | 7,738,607 |
| Continuous | Low | CNR | 19088 | 2064 | 97292 | -19 | -1 | -1 | 3.34 (2.32%) | 162,359 | 1,176,433 |
|  |  | CNR & responsible ownership | 16836 | 1701 | 98392 | -29 | -18 | 0 | **3.36 (3.01%)** | 157,107 | 1,000,141 |
|  |  | 0.5 CNR & 0.5 sheltering | 17971 | 5722 | 98720 | -24 | 174 | 0 | 3.31 (1.70%) | 366,744 | 3,131,743 |
|  |  | **Culling** | **16229** | 2071 | 97612 | **-31** | -1 | -1 | 3.32 (1.84%) | 130,634 | 1,147,611 |
|  |  | Responsible ownership | 21739 | 1719 | 99475 | -8 | -18 | 1 | 3.28 (0.57%) | **124,844** | **976,036** |
|  |  | Sheltering | 16436 | 10450 | 100553 | -31 | 401 | 2 | 3.29 (0.87%) | 611,772 | 5,588,233 |
|  | Medium | CNR | 16922 | 2053 | 96786 | -28 | -2 | -2 | 3.37 (3.48%) | 172,574 | 1,181,859 |
|  |  |  | S | H | O | S | H | O |  | Shelter staff-to-dog ratio |  |
|  |  |  |  |  |  |  |  |  |  | High | Low |
|  |  | **CNR & responsible ownership** | **10277** | 1445 | 99071 | **-57** | -31 | 1 | **3.42 (4.94%)** | 150,133 | 870,750 |
|  |  | 0.5 CNR & 0.5 sheltering | 14607 | 6722 | 98564 | -38 | 222 | 0 | 3.35 (2.80%) | 435,248 | 3,736,621 |
|  |  | Culling | 10917 | 2059 | 97072 | -54 | -1 | -1 | 3.37 (3.32%) | 132,644 | 1,144,582 |
|  |  | Responsible ownership | 19466 | 1466 | 100547 | -18 | -30 | 2 | 3.30 (1.22%) | **120,863** | **851,095** |
|  |  | Sheltering | 11193 | 13438 | 101065 | -53 | 544 | 3 | 3.32 (1.80%) | 799,722 | 7,300,451 |
|  | High | CNR | 15177 | 2046 | 96461 | -36 | -2 | -2 | 3.40 (4.28%) | 176,965 | 1,183,033 |
|  |  | **CNR & responsible ownership** | **2477** | 1261 | 99994 | **-90** | -40 | 2 | **3.48 (6.71%)** | 130,727 | 762,187 |
|  |  | 0.5 CNR & 0.5 sheltering | 11571 | 7034 | 98320 | -51 | 237 | 0 | 3.38 (3.70%) | 460,738 | 3,969,868 |
|  |  | Culling | 6445 | 2049 | 96612 | -73 | -2 | -2 | 3.41 (4.70%) | 132,734 | 1,140,277 |
|  |  | Responsible ownership | 16560 | 1281 | 101555 | -30 | -39 | 3 | 3.33 (2.04%) | **120,690** | **761,314** |
|  |  | Sheltering | 6674 | 13929 | 100805 | -72 | 568 | 2 | 3.35 (2.89%) | 849,490 | 7,753,805 |

\* As annual interventions did not reach a single stable equilibrium point but varied between two points due to the annual interventions, the average of the minimum and maximum equilibrium population size was reported.

The equilibrium population size fluctuated between annually applied interventions. Annual interventions that targeted removal from the population (i.e. culling and sheltering) fluctuated more than those that targeted recruitment (i.e. responsible ownership and CNR). When comparing the continuously applied interventions with the average of the minimum and maximum equilibrium population size for annual interventions, the resulting equilibrium population sizes were almost equal. Overall, the greatest proportion of neutered dogs was 0.77, achieved by annually applied high coverage combined CNR and responsible ownership (Figure 3).

**Figure 3.**
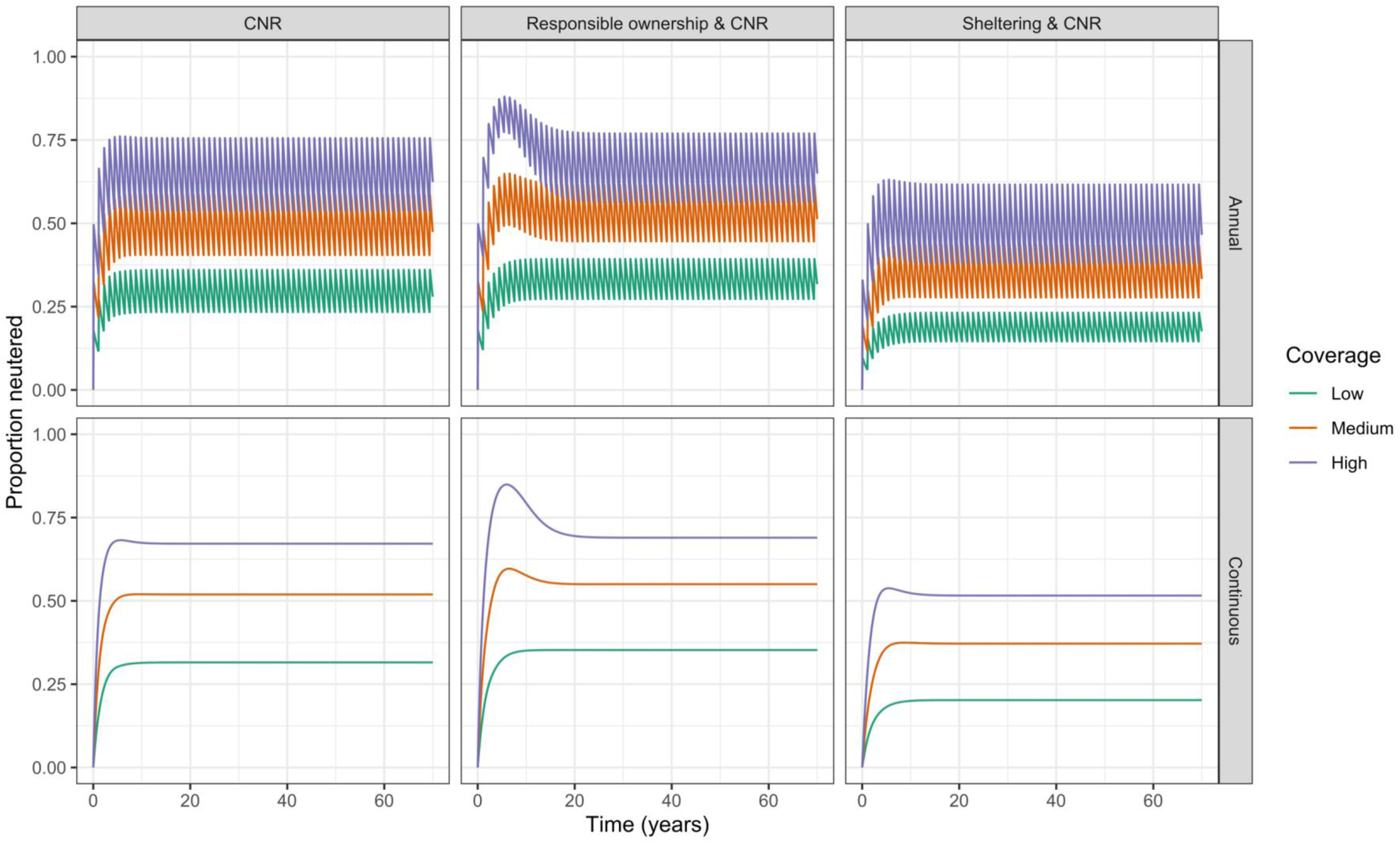
Proportion of street dogs neutered for interventions including CNR and applied for duration of the simulation at low, medium, and high coverages applied annually and continuously.

When applied continuously at a high coverage, responsible ownership was the cheapest intervention (average annual costs €120,690 for high, and €761,314 for low staff-to-dog ratio; Table 4), but only slightly cheaper than combined CNR and responsible ownership (€130,727 for high, and €762,187 for low staff-to-dog ratio). Combined CNR and responsible ownership was cheaper than CNR applied alone. This is due to the effectiveness of the combined method at reducing street dog population size, resulting in fewer dogs being neutered to maintain the neutering coverage. Sheltering applied continuously at high coverages was most expensive (€849,490 for high, and €7,753,805 for low staff-to-dog ratio). For interventions applied annually (responsible ownership was not included as continuous effects were assumed), culling was the cheapest intervention for all coverages for high staff-to-dog ratios, and combined CNR and responsible ownership was the cheapest intervention for all coverages for low staff-to-dog ratios.

The greatest improvement in overall welfare score was achieved by combined CNR and responsible ownership for interventions applied continuously and annually at all coverages (Table 4), improving the welfare score by 6.58% and 6.71% respectively. The lowest improvement in welfare score was by responsible ownership at low coverages, which improved the welfare score by 0.57%.

Results of the global sensitivity analysis for the impact of interventions on each subpopulation are presented in Figures S5-10. The simulated street dog populations stabilised at an equilibrium between 12,823 and 18,619 for CNR; 4,548 and 13,599 for culling; 4,530 and 14,248 for sheltering; 7,736 and 23,539 for responsible ownership; 1,496 and 10,354 for CNR and responsible ownership; and 8,750 and 17,212 for combined CNR and sheltering.

### Results for interventions applied for a single five-year period

Table 5 outlines the results for interventions applied for five years. For interventions applied annually and continuously, culling achieved the greatest reduction in street dog population size at all coverages (low: annual 34%, continuous 29%; medium: annual 57%, continuous 51%; high: annual 77%, continuous 71%), though sheltering was almost equally effective (Figure 2).

**Table 5.**
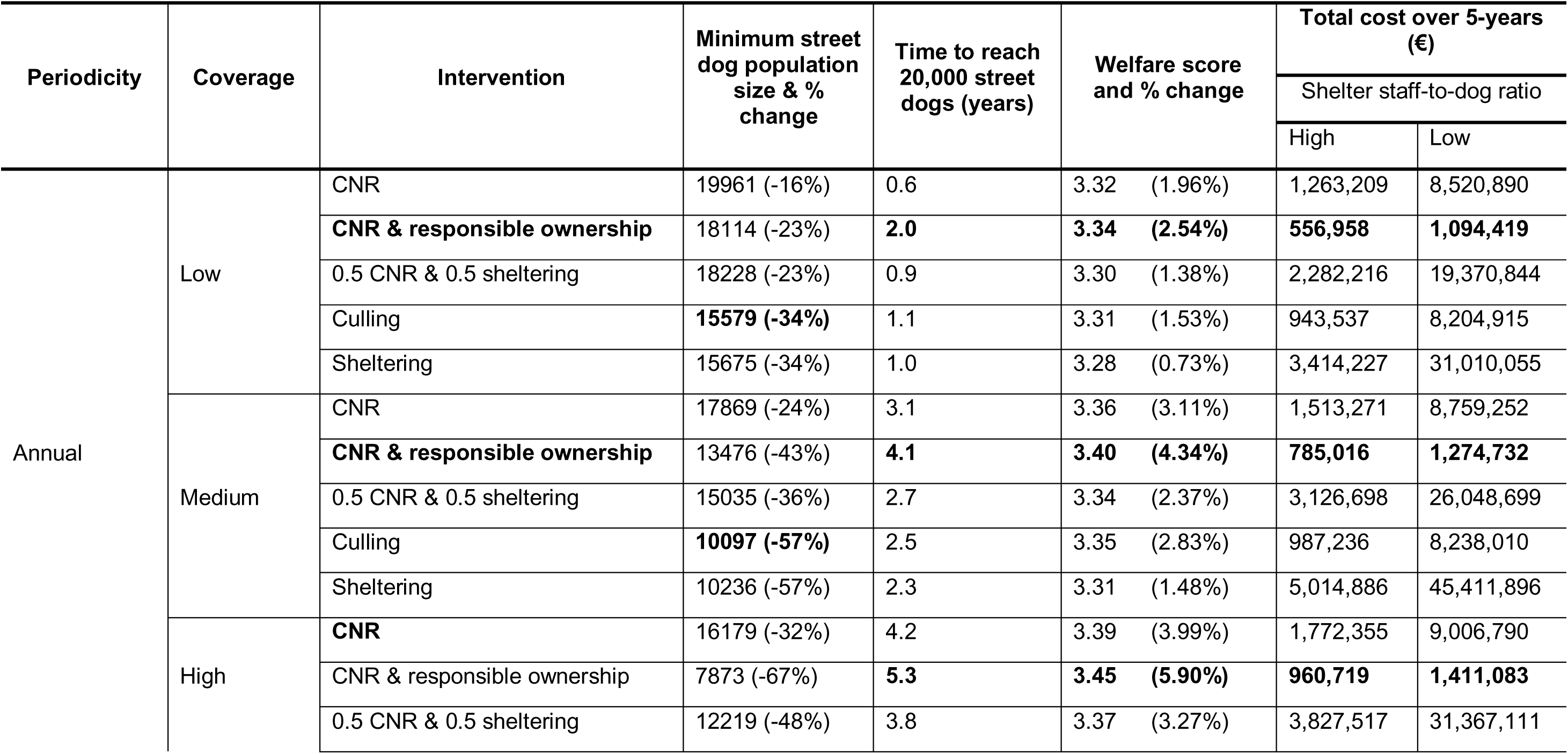
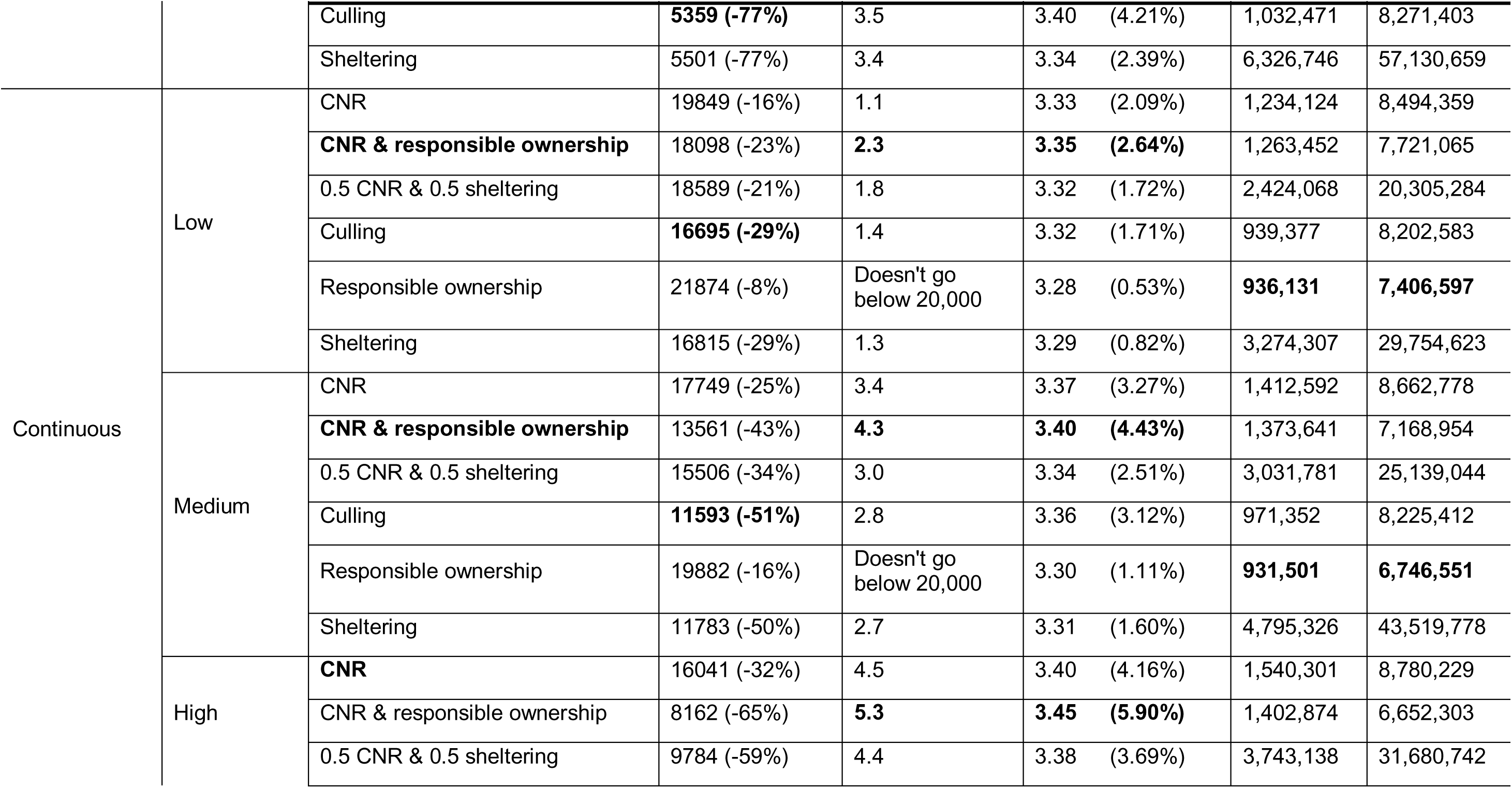
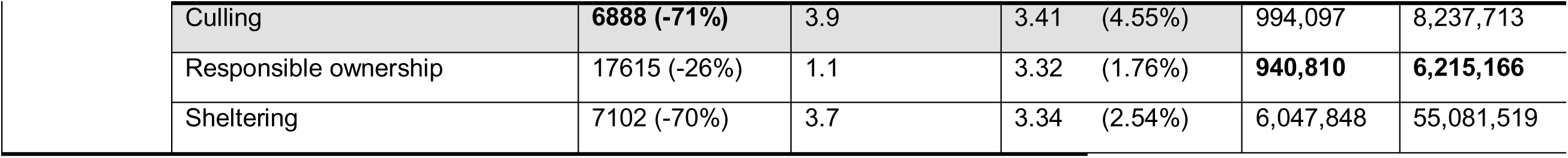
Impact of five-year intervention on minimum street dog population size and time taken between end of intervention and street dog population size reaching 20,000 dogs. The intervention with the overall longest time to reach baseline population size is highlighted in grey. The intervention with the greatest reduction in street dog population size, the longest time to reach baseline population size, the greatest increase in welfare score, and the lowest cost for each periodicity-coverage combination are highlighted in bold.

The street dog population returned most slowly to above 20,000 dogs in combined CNR and responsible ownership interventions for all coverages and periodicities (Table 5). For example, at high continuous coverages, the street dog population reached above 20,000 dogs after 5.3 years, compared to 1.1, 3.7, and 3.9 years for responsible ownership, sheltering and culling respectively. The street dog population returned more slowly to above 20,000 dogs for interventions with high coverages (i.e. due to higher magnitudes of effect). There was little difference between annual and continuous interventions.

The intervention with the lowest total cost over the 5-year intervention was annually applied combined CNR and responsible ownership at a low coverage (€556,958 at high and €1,094,419 at low staff-to-dog ratios; Table 5). The most expensive intervention was sheltering applied annually at a high coverage (€6,326,746 at high and €57,130,659 at low staff-to-dog ratios).

The greatest improvement in overall welfare scores for both annual and continuous interventions was achieved by combined CNR and responsible ownership when applied at a high coverage (increase of 5.90% for both annual and continuous; Table 5). Responsible ownership applied continuously had the least improvement in welfare score when applied at a low coverage. This reduced the welfare score by 0.53%.

## Discussion

This study provides important information on the predicted effectiveness of dog population management interventions whilst (i) explicitly incorporating the interactions between the different subpopulations, (ii) modelling a range of management methods at different periodicities and coverages, and (iii) also providing estimates of the welfare impacts and costs. Our results indicate that methods targeting multiple sources of population increase, such as combined CNR and responsible ownership interventions, may be most effective at reducing street dog population size when applied over longer periods of time, whilst also remaining cost-effective and maintaining highest levels of overall dog welfare. Sheltering and culling may be more effective over shorter periods of time but, compared to methods including neutering, the population may rapidly return to an equilibrium population size once management has ended. This model provides a template for assessing the impact of free-roaming dog population management in multiple contexts (e.g. rural, semi-rural, or urban populations) by applying population-specific parameters.

Reducing free-roaming dog population size can reduce the risks associated with free-roaming dogs (e.g. public health, livestock production, wildlife conservation). The results of this study suggest that combined CNR and responsible ownership may most greatly reduce these risks. Combined CNR and responsible ownership had a synergistic effect: CNR and responsible ownership applied in isolation were least effective at reducing street dog population size. These findings are similar to those reported in previous modelling studies that also suggest a synergistic effect when combining fertility control with reduced owned dog abandonment ^[15,23]^. The synergistic effect can be explained by this method reducing multiple flows that can replenish the population size; without targeting multiple flows, the potential for the population to increase in size remains. Targeting multiple flows by combining responsible ownership interventions with other methods, such as culling or sheltering, may also have had strong reductive effects on the street dog population size. For example, combined culling with responsible ownership would increase the street dog death rate and reduce the flow of abandoned owned dogs to the street dog population. Indeed, other combinations of methods may also have been effective, such as combined CNR and culling. Though it is important to consider the logistical challenges in combining different methods. For example, combined CNR and culling may be challenging to apply in reality, as neutered dogs would need to be recognised and not culled to ensure efforts were not wasted. In this study, we only investigated methods that would feasibly be applied and have often been reported ^[27]^.

When considering the individually applied interventions, culling and sheltering were most effective at reducing the street dog population size at all intervention coverages and periodicities (Figure 2). Both culling and sheltering increase the rate of removal from the street dog population; culling by increasing the death rate and sheltering by increasing the sheltering rate. CNR and responsible ownership both reduce recruitment to the street dog population; CNR by reducing birth rates and responsible ownership by reducing owned dog abandonment rates. Our results suggest that, when considering interventions applied in isolation, increasing removal rates are more effective at reducing street dog population sizes than reducing recruitment rates. Our findings are similar to those reported by Amaku *et al*., (2010) ^[17]^, who also reported culling to be more effective than CNR applied alone, but contradict those reported by Yoak *et al*., (2016) ^[25]^ and Hogasen *et al*., (2013) ^[22]^ who predicted CNR to be more effective than culling or sheltering at reducing street dog population size. This difference may be due to the differences in mathematical modelling methods and assumptions. Yoak *et al*., (2016) ^[25]^ implemented an agent-based modelling approach, compared to the differential equations applied by Amaku *et al*., (2010) and in our study.

Although our results suggest that CNR applied alone would be one of the least effective methods, there has been increasing interest over the last 20 years in the use of CNR to manage free-roaming dog populations ^[27]^. Empirical studies have reported CNR to be effective in reducing population sizes between 12% over 1.5 years and 40% over 12 years at varying reported intervention coverages ^[18,44–46]^. Our results suggest CNR may reduce the street dog population size up to 36% (high coverage) over 5 years, though it is hard to compare these results to empirical studies due to a lack of reporting of intervention coverage and management length (see ^[27]^ for review).

There has been little focus on the impact and effectiveness of responsible ownership campaigns on street dog population size or human behaviour change ^[27]^. Our results suggest that targeting efforts to reduce abandonment could be as important as directing efforts to reduce the birth rate or increase the removal rate. These findings are in line with those in previous modelling studies reporting synergistic effects of combined fertility control and restricted movement of owned dogs ^[26]^, and emphasising the potential impact of owned dog abandonment dampening the effectiveness of CNR interventions ^[15,17,22]^. Future dog population management efforts should aim to identify the sources of population increase (i.e. abandonment or immigration) to understand the potential impact of CNR and other interventions.

Despite the promising potential effects of combined CNR and responsible ownership, the evidence is lacking on whether responsible ownership interventions can actually decrease abandonment rates and increase shelter adoption rates to the degree explored in this study. Our lack of understanding of the effect of responsible ownership campaigns may be due to challenges in quantifying dog ownership practices. It is particularly challenging to accurately quantify the rate of owned dog abandonment, as this is likely to be under-reported ^[34]^. Future studies may consider quantifying abandonment through questionnaire surveys, focus groups, or using local shelter relinquishment figures as an indicator of abandonment. It is important to measure the impact of responsible ownership campaigns, in combination with CNR efforts, to identify the most effective strategies in reducing abandonment rates and increasing responsible ownership practices.

In simulations where interventions were applied for five years, culling was most effective at reducing the street dog population size at all management coverages. In the present study, both CNR and combined CNR and responsible ownership were more effective at maintaining the street dog population at a lower size for longer than other interventions in the five-year intervention simulation (i.e. it took longer for the population to reach above 20,000 dogs after management ended). Individuals that have been removed through culling or sheltering may quickly be replaced through births or immigration, resulting in the population quickly returning to the carrying capacity. The longer lasting effects of neutering may be explained by the neutered individuals remaining in the population and, as they do not contribute to birth, dampening the potential for population growth.

Unsurprisingly, interventions including sheltering were most expensive overall (Table 4 and Table 5). Sheltering increases the shelter dog population size and, if the rehoming rates stay the same, greater staff resources are required to care for the shelter dog population. Killing of shelter dogs would reduce the shelter dog population and thus reduce costs, but we did not model this as several countries have a no kill policy for shelter dogs ^[47]^, only allowing euthanasia for behavioural or health problems. In these countries, sheltering as a method of population control may result in an increase in the shelter dog population and, without an improvement in rehoming rates, has the potential to lead to life-long stays in shelters and overcrowding. As this method is costly, it is potentially only a feasible option for higher income countries ^[10]^.

The cheapest intervention overall in terms of staff costs was responsible ownership campaigns. Responsible ownership campaigns do not require dog catchers, veterinarians, or veterinary nurses. Although when applied alone this method was the least effective, the combination of CNR and responsible ownership was more effective at reducing the population size and was only marginally more costly than responsible ownership alone. As this method was effective in reducing population size, fewer dog catchers, veterinarians and nurses were required throughout the simulation, making it more cost-efficient in the long-term. Culling and CNR were more similar in costs, in terms of staff resources, than might be anticipated (Table 4 and Table 5). Whilst CNR requires higher-paid workers (veterinarians and veterinary nurses), as the simulation progressed, there were fewer intact dogs to neuter and fewer veterinarians were required to maintain the intervention coverage. Culling required a higher number of dog catchers to maintain the intervention coverage throughout the simulation, resulting in there being less of a difference between culling and CNR in terms of staff resources needed.

Our study has limitations in the cost analysis, as this did not encompass the full cost of the interventions, as it excluded staff training (e.g. in catching dogs, performing surgeries), facilities (e.g. clinics, kennels) and equipment costs. For example, creating and buying advertising space to promote a responsible ownership campaign may cost tens of thousands of euros (VIER PFOTEN International, personal communication), and this is not incorporated into the analysis. The cost of management is rarely provided by studies investigating the impact of dog population management ^[27]^. Where costs do exist, they exist in disparate datasets: differing by country (i.e. country specific costs) and methods of application. For example, for CNR, costs may be lower for interventions that involve less staff-training and shorter pre- and post-surgical holding times, but may compromise dog health and welfare as a result ^[48]^. Similarly, costs of running a shelter will vary depending on the shelter environment and management: shelters with larger dog-pen sizes, less enrichment, and lower staff-to-dog ratios may be less costly, but may also compromise health and welfare ^[49]^. There are currently no data available to quantify the costs required for a responsible ownership campaign to reduce the abandonment of owned dogs by a quantified amount. To fully assess and compare the costs of different interventions requires integration of multiscale datasets across the dog population management field. Whilst this study does not encompass the full costs of interventions, calculating the staff-time costs required allowed us to evaluate intervention costs on a comparable scale.

In addition to population size, welfare measures are important indicators of dog population management impact ^[11]^. The greatest improvement in welfare score, compared to the baseline simulation, was achieved by combined CNR and responsible ownership. This is due to an overall reduction in street dog population size and an increase in the proportion of street dogs that were neutered, whose welfare is rated more highly than intact street dogs. However, as we used aggregated welfare scores, weighted by self-reported knowledge of dog subpopulations provided by veterinarians in Italy ^[11]^, the scores may not be applicable to all countries. For example, the welfare of the street dog population in one country may be greater than another, due to country-specific risks to dog health and welfare. It is challenging to compare the welfare impact due to a lack of comparable welfare data within and between subpopulations ^[27]^. Using this overall welfare score allows us to compare the potential welfare impact of different interventions on overall welfare.

Dog population management methods may also have a short-term impact on dog health and welfare, depending on the method used. For example, there are important welfare risks related to CNR interventions, depending on dog handling procedures, standards of surgery, and post-operative care ^[48,50]^. The impact of culling on dog welfare also depends on the method used. Historically, culling methods have been inhumane ^[51,52]^. In some locations, mass killing is carried out by distributing poisons, most commonly strychnine ^[51]^ which is not a recommended method of killing from an animal welfare perspective ^[10]^. In locations where poisoning is not used, the methods to restrain the dogs may also be of welfare concern. For example, methods of physical restraint include the holding of body parts by rope or metal tongs which can cause laceration or tissue damage ^[51]^. Once restrained, methods to kill dogs have included electrocution, carbon monoxide poisoning and drowning ^[51]^. These methods are not recommended by the World Organisation for Animal Health (OIE) but are still employed in some areas ^[10]^. These additional welfare concerns should be considered when determining appropriate dog population management intervention.

Several assumptions were made by this model. Firstly, the model did not differentiate between different age and sex categories. All individuals were assumed to contribute equally to the dynamics occurring between the different subpopulations. Currently the data is lacking at the level necessary to model the flows between subpopulations for different age and sex categories. Further study to determine these rates would be beneficial for informing future understanding of the dynamics between dog subpopulations. The model also did not include the effects of immigration and emigration explicitly. If interventions occur in neighbourhoods, instead of homogeneously throughout a city, there is the potential that this could increase movement of dogs from other parts of the city. We assumed that methods could be applied at consistent levels of coverage, regardless of the street dog population size. In reality, it may be more challenging to find dogs when the street dog population size is smaller.

The results of this study primarily apply to free-roaming dog populations in Europe. Populations in other geographic locations may differ in their movement within and between the subpopulations. Cultural factors within countries, such as religious beliefs may affect public attitudes and acceptance of management methods. In areas with lower abandonment rates and/or opposition to neutering, combined CNR and responsible ownership interventions may be less effective. Despite this, this model highlights the importance of identifying sources of population increase and can be used by applying population specific parameters to identify effective and efficient management methods.

## Conclusions

System dynamics modelling is a useful tool for investigating the potential impact of different dog population management methods. Overall, combined CNR and responsible ownership may be the most effective at reducing street dog population size, maintaining the population at a lower size, whilst also improving the overall welfare score of dogs, at low costs in terms of staff resources. Future dog population management would benefit from identifying and targeting all potential flows that may increase the street dog population size (such as abandonment and immigration). More studies are required to measure the impact of interventions, such as responsible ownership campaigns, on human behaviour change.

## Data availability

Data available from https://github.com/lauren-smith-r/Smith-et-al-dog-pop-management-systems-modelling.

## Acknowledgements and funding sources

We are grateful to VIER PFOTEN International for funding this project and Dr Paolo Dalla Villa for assisting in supervision of the project. We also thank Sarah Ross for providing important contacts and information to assist in the study.

## Author contributions

L.M.S., L.M.C., R.J.Q., A.M.M., and S.H. conceptualised the study; L.M.S., L.M.C., R.J.Q. designed research; L.M.S. collected data; L.M.S., R.J.Q., C.G., and L.M.C. analysed and interpreted the data; L.M.C. and R.J.Q supervised the project; L.M.S. wrote the manuscript; L.M.S., R.J.Q., C.G., A.M.M., S.H., and L.M.C. reviewed the manuscript.

## Competing interest statement

The authors declare that: A.M.M. and S.H. are employed by VIER PFOTEN International, an animal welfare organisation; L.M.C has received a research grant from VIER PFOTEN International; and L.M.S.’s PhD research has been funded by VIER PFOTEN International.

## Notes

https://github.com/lauren-smith-r/Smith-et-al-dog-pop-management-systems-modelling

